# Test-Retest Reliability of Task-based and Resting-state Blood Oxygen Level Dependence and Cerebral Blood Flow measures

**DOI:** 10.1101/381251

**Authors:** Štefan Holiga, Fabio Sambataro, Cécile Luzy, Gérard Greig, Neena Sarkar, Remco J Renken, Jan-Bernard C Marsman, Scott A Schobel, Alessandro Bertolino, Juergen Dukart

## Abstract

Despite their wide-spread use, only limited information is available on the comparative test-retest reliability of task-based functional and resting state magnetic resonance imaging measures of blood oxygen level dependence (tb-fMRI and rs-fMRI) and cerebral blood flow (CBF) using arterial spin labeling. This information is critical to designing properly powered longitudinal studies. Here we comprehensively quantified and compared the test-retest reliability and reproducibility performance of 8 commonly applied fMRI tasks, 6 rs-fMRI metrics and CBF in 30 healthy volunteers. We find large variability in test-retest reliability performance across the different tb-fMRI paradigms and rs-fMRI metrics, ranging from poor to excellent. A larger extent of activation in tb-fMRI is linked to higher between-subject reliability of the respective task suggesting that differences in the amount of activation may be used as a first reliability estimate of novel tb-fMRI paradigms. For rs-fMRI, a good reliability of local activity estimates is paralleled by poor performance of global connectivity metrics. Evaluated CBF measures provide in general a good to excellent test-reliability matching or surpassing the best performing tb-fMRI and rs-fMRI metrics. This comprehensive effort allows for direct comparisons of test-retest reliability between the evaluated MRI domains and measures to aid the design of future tb-fMRI, rs-fMRI and CBF studies.

## Introduction

Functional magnetic resonance imaging (fMRI) based sequences such as task-based, resting-state blood oxygenation level-dependent MRI (BOLD; tb-fMRI and rs-fMRI) and arterial spin labelling (ASL) of regional cerebral blood flow (CBF) are now commonly applied for studying human brain function [1–7]. Beside their widespread application in systems neuroscience, they are also recognized as valuable indices for investigating aberrant neural mechanisms behind a variety of psychiatric and neurological diseases and for evaluation of experimental interventions [8–11]. In particular, their application as diagnostic, stratification, pharmacodynamic and efficacy biomarkers has been suggested in that context [8, 12–14].

Various derived measures ranging from local activity estimates to local and global connectivity metrics have been suggested for all of the above MRI measures [4,15,16]. Given the complementary nature of tb-fMRI, rs-fMRI and CBF measures their combined acquisition may provide better insights into understanding of underlying pathophysiological processes and potential treatment effects. In addition to this sensitivity to relevant disease or treatment-induced alterations, an important criterion for selection and integration of MRI measures into clinical studies is also their reliability in a longitudinal setting.

Test-retest reliabilities of the aforementioned MRI measures have been extensively evaluated, with strongly varying reliability estimates ranging from poor to excellent [17–25]. Despite this extensive research, longitudinal consistency of tb-fMRI, rs-fMRI and CBF measures was typically established in separate studies, using different hardware, pre-processing and analysis methodology. Furthermore, studies performing comparisons of different metrics extracted from those fMRI data mainly focused on within domain evaluation, i.e. by comparing different rs-fMRI metrics. Therefore, little is known about the *relative* reliabilities of these measures. The methodological discrepancies consequently limit comparability of reliability estimates for different MRI domains across studies [20,24].

Here, we addressed these limitations by conducting a comprehensive dedicated methodological study comparing 8 established fMRI tasks covering various neuropsychological domains, 6 established rs-fMRI metrics and quantitative CBF evaluated in the same subjects using the same hardware, preprocessing and analysis methodology.

## Materials and Methods

### Study population and Criteria for Inclusion

Thirty one healthy male and female subjects (Age: 25 ± 5 years [mean ± standard deviation]; 7 males/24 females) participated in the study after providing written informed consent..

Health status was determined by screening assessments and principal investigator judgment and was defined by the absence of any active or chronic disease or positive signs on a complete physical examination including vital signs, 12-lead electrocardiogram, hematology, blood chemistry, serology and urinalysis. Only subjects with a body mass index (BMI) between 18 to 30 kg/m^2^ with a body weight between 50-100 kg were included in the study. All subjects were fluent in the language of the investigator and were able to comply with study requirements as judged by the principal investigator.

The study was carried out according to local regulations and the International Council for Harmonisation of Technical Requirements for Pharmaceuticals for Human Use (ICH) guidelines. All experimental procedures conformed to the Declaration of Helsinki and the study protocol was approved by the local ethics committee (Foundation Beoordeling Ethiek Biomedisch Onderzoek, Assen, Netherlands; fMRI - RHE323EC-153231 - NL54292.056.15). The study has been registered on Clinicaltrials.gov under the identifier NCT02560142 and was sponsored by F. Hoffmann-La Roche Ltd.

### Study Design

All study visits were performed at a single center (Neuroimaging Center, University Medical Center Groningen, Netherlands). A screening period of 28 days (15±3 days before the baseline Visit 1) preceded the study assessment period. Subsequently, two study visits (Visit 1; Visit 2) were performed fourteen days apart. The imaging protocol consisted of a series of structural and functional MRI sequences/tasks, as outlined in Table S1.

The battery of structural MRI (see Table S1, MRI measures 1-3) was performed for visualization and data processing purposes, and to rule out incidental neuroradiological findings. During screening, the subjects were trained in the completion of all tb-fMRI tasks using training versions of the tasks. Only subjects with adequate performance were included in the study. A resting scanning session at screening (Table S1, MRI measures 4-5) was added to in order to minimize the magnitude of the putative habituation effect between Visit 1 and Visit 2. The order of the structural and rs-fMRI measures (Table S1, MRI measures 1-5) was fixed for each subject and visit. The order of the tb-fMRI in Visit 1 and Visit 2 (Table S1, MRI measures 6-10) was randomized across subjects using the Williams (Latin Squares) design [26] to account for potential carry-over effects. Right before the start of the particular imaging session or task, the participant received an operator-guided on-screen reminder to reassure understanding of the particular task and all associated procedures.

### MRI Data acquisition

All scans were performed by experienced MRI technicians on a 3 T clinical scanner (Intera, Philips Healthcare, Best, Netherlands) using a 32-channel head coil. *T_1_*-weighted images were obtained using a 3D fast field echo (FFE) sequence (repetition time, *TR* = 10.4 ms; echo time, *TE* = 5.7 ms; flip angle, *FA* = 8°; 160 slices; in-plane resolution = 1 × 1 mm^2^; slice thickness 1 mm). For CBF computation 60 pairs of labeled and control images with 17 axial slices, 7 mm slice thickness and no gap covering the whole brain were collected using a pseudo-continuous arterial spin labeling (pCASL) sequence (*TR* = 4000 ms; *TE* = 14 ms, *FA* = 90°; labeling duration = 1650 ms; post-labeling delay = 1600 ms; labeling gap = 2cm; in-plane resolution = 3 × 3 mm^2^). A 2D single-shot echo-planar imaging (EPI) readout with fat suppression was used. Additionally, a separate proton density image (*M_0_*) was collected to obtain voxel-wise intensity of fully relaxed blood spins. For rs-fMRI, 244 volumes of BOLD effect sensitive images covering the whole brain were acquired using a gradient-echo EPI sequence (*TR* = 2000 ms, *TE* = 30 ms; *FA* = 90°; 39 axial slices with 1 mm gap, nominal in-plane resolution 3 × 3 mm^2^; slice thickness at 3 mm). The same EPI sequence and the same imaging parameters except the number of volumes were used to acquire BOLD signal during performance of the respective tb-fMRI tasks.

### MRI preprocessing and analyses

All preprocessing and statistical analyses were performed using Matlab (R2013b, The MathWorks Inc., Natick, MA, USA) and SPM12 (Wellcome Trust Centre for Neuroimaging, UCL, London, UK). Quantitative CBF maps were computed according to recommendation of the ISMRM Perfusion Study Group and the European Consortium for ASL in Dementia [27]. Preprocessing of CBF and BOLD data comprised motion correction, distortion correction (for BOLD), spatial registration to a structural scan with a subsequent normalization into the Montreal Neurological Institute (MNI) space, masking of non-grey matter voxels and smoothing with a Gaussian kernel of 6 mm full-width at half maximum.

### rs-fMRI measures

Motion was regressed out of the rs-fMRI data using the Friston 24-parameter model approach alongside with mean white matter and CSF signal [28,29]. The following rs-fMRI measures were calculated: degree centrality (DC), fractional and absolute amplitude of low frequency fluctuations (fALFF and ALFF, respectively), regional homogeneity (ReHo), eigenvector centrality (EC) and Hurst exponent. ALFF, fALFF, ReHo (coherence), and DC were computed using the Rest toolkit [15], EC as implemented by Wink et al. [30], and Hurst exponent as implemented by Maxim et al. [31]. In brief, DC is a count-based measure that assigns to each voxel the sum of all correlation coefficients between the time series of that voxel and all other voxels in the brain exceeding a prespecified threshold (r>0.25). A recommended correlation threshold of 0.25 was used for DC computation to eliminate counting voxels that have low temporal correlation attributable to signal noise [32]. DC maps were additionally *z*-transformed to reduce the effects of global connectivity changes. Temporally unfiltered time series were used for estimation of Hurst exponent. ALFF and fALFF reflect the absolute and normalized amplitude of local temporal low frequency fluctuations. ReHo represents the coherence of a voxels time series with its immediate neighborhood. EC represents the importance of a voxel in a network based on its synchronization strength to other more or less important regions. Hurst exponent provides a measurement of persistence of specific signals in the time series. All measures were computed as suggested in the respective cited publications using default parameters, including removal of a linear trend and restriction to the low frequency range (for fALFF divided by the amplitude of frequencies outside the range) used by the REST toolkit (0.01–0.08 Hz) [15].

### tb-fMRI measures

Details of the employed experimental paradigms are summarized in the Supplementary Material. In brief, the following established fMRI paradigms were evaluated: reward expectation – monetary incentive delay task (MID), working memory – N-back task, theory of mind – ToM, emotional face matching – FM, response inhibition – Go/No-go, memory encoding, recall and recognition. To determine task-dependent activation, (first-level) *t*-contrasts of ‘active vs control’ condition were computed for each fMRI task per subject and session (Face matching: Faces > Shapes, MID: Win > Control, N-back: 2 back > 0 back, Go/No-go: No-go > Go, Encoding, Recall and Recognition: Professions > Ears, ToM: Affective > Visuo-spatial). Effects of motion were controlled for in all tasks by including 6 motion parameters (translation and rotation) in all models. Group-level main effects of task ((de)activation maps) were evaluated using the obtained individual contrast maps for all fMRI tasks including estimates for all subjects and visits in a voxel-wise manner using a family-wise error (FWE) corrected threshold of *p*<0.05. Additionally, separate group (de)activation tests were computed for the two visits.

### Reliability analyses

To evaluate the reliability of respective tb-fMRI, rs-fMRI and CBF measures, we computed two types of intra-class correlation (ICC) [33] and consistency metrics as described below. For ICCs the following criteria as developed by Cicchetti and Sparrow [34] were applied for interpretation: poor (below 0.4), fair (0.4–-0.59), good (0.6–-0.74), and excellent (≥0.75). Two types of ICCs were used for all analyses: ICC(2,1) for directly derived measures (i.e. % correct or voxel-wise activation) and ICC(2,k) for average measures from the respective visits (i.e. reaction times or average activation from regions-of-interest) [33]. Specifically, for ICC(2,1), a two-way random effects model (column and row effects random) was used to calculate the degree of consistency among measurements. This model is also known as norm-referenced reliability and as Winer’s adjustment for anchor points. For ICC(2,k), a two-way random effects model (column and row effects random) was used to calculate the degree of absolute agreement for measurements that are averages based on *k* independent measurements on randomly selected objects [35].

Importantly, for both types of ICCs a maximum possible positive value of 1 indicates perfect reliability. In contrast, the applied ICCs are not limited in terms of their lower bound with negative coefficients below -1 being possible in case of anti-correlation.

### Behavioral reliability analyses

First, we evaluated the stability and reliability of behavioral measures acquired during the fMRI tasks. For this we computed paired *t*-tests evaluating changes in mean performance across visits for the respective measures. Further, we assessed the test-retest reliability of behavioral measures acquired during the specific fMRI tasks using the ICCs described above.

### Voxel-wise reliability analyses

To estimate the reliability of the various tb-fMRI, rs-fMRI and CBF measures we computed several types of reliability and consistency estimates to estimate voxel- and region-wise reliability and consistency of the respective measures. To characterize the consistency of activation patterns observed with fMRI at session 1 and 2, we computed Jaccard indices of overlap of the whole-brain activity between visit 1 and visit 2 (area of overlap divided by the overall activated area), by systematically varying the cut-off activation threshold (*t*-value) for both visits and counting concordant/discordant pairs of (de)activated voxels. Further, to characterize the voxel-wise test-retest reliability of the respective measures we computed voxel-wise intra class correlation coefficients (ICC(2,1)) for all tb-fMRI contrast maps, rs-fMRI and CBF measures. As 3 visits were available for rs-fMRI and CBF, voxel-wise ICCs were computed between screening and visit 1 and between visit 1 and visit 2 (consistent with fMRI tasks). Median ICCs of all significantly (de)activated voxels were then extracted from the obtained voxel-wise ICC maps. Further, median voxel-wise ICCs were extracted for rs-fMRI and CBF from pre-specified, commonly used resting state networks (http://findlab.stanford.edu/functional_ROIs.html). For tb-fMRI, median voxel-wise ICCs were computed separately within regions showing significant task-related activation or deactivation (pooled over both visits). Additionally, to evaluate the consistency of the average voxel-wise group activation maps obtained at visits 1 and 2, we computed test retest reliability (ICC(2,k)) between the spatial activation profiles obtained at both visits (*t*-contrasts). Lastly, we aimed to evaluate if the amount of observed task-induced activation or deactivation was linked to the respective test-retest reliability. For this we computed a Pearson correlation between both visits across all tasks.

### Region-wise reliability analyses

We further aimed to evaluate if averaging over specific brain regions affected the reliability estimates. Region-wise ICCs were computed for all measures by extracting mean values from regions provided by the automated anatomical labeling (AAL) atlas (116 regions). Within-region/between-subject and between-region/within-subject ICCs (ICC(2,k)) were computed for each region and each imaging measure to evaluate the between-subject and within-subject reliabilities, respectively. The first type of ICCs (within-region/between-subject) thereby reflects the reliability of the signal within a specific a region across subjects (i.e. where the order of subjects remains the same). The second type (between-region/within subject) provides an estimate of the robustness of the observed spatial activation pattern within each subject (i.e. does region A show a consistently higher activation as compared to region B?). For rs-fMRI and CBF, all ICCs were computed for screening to visit 1 and for visit 1 to visit 2. The ICCs related to tb-fMRI were calculated for visit 1 to visit 2. Reliability of the mean (de)activation within significant regions was also evaluated for tb-fMRI data. Additionally, as specific regions are of particular interest for some of the included tasks (ventral striatum for MID, dorsolateral prefrontal cortex for N-back, left and right amygdala for FM and medial prefrontal cortex for ToM) test-retest reliability (ICC(2,k)) was computed for mean activation values extracted from these regions (defined using corresponding anatomical clusters showing significant activation at both visits).

## Results

### Obtained data

All subjects complied with the study protocol and finished the required assessments. One subject (ID 1207) was excluded from the study due to a newly diagnosed attention deficit hyperactivity disorder. All participants were able to perform the fMRI tasks. Based on quality check, the CBF scans for one subject, rs-fMRI data for 2 subjects and N-back data for 1 subject were discarded due to insufficient coverage due to misplaced bounding box and/or excessive motion. Overall, this resulted in evaluable data for 28 to 30 subjects depending on the respective fMRI domain.

### Results of behavioral reliability analyses

Mean reaction times significantly decreased at visit 2 for the MID, FM, Encoding and the 0-back condition of the N-back task (Table S2). The number of hits and the collected reward significantly increased in the MID task. No differences were observed for other behavioral indices for any of the tasks except a slight increase in the miss rate for the 2-back condition of the N-back task. Reaction times in the control conditions of all tasks except Go/No-go and recognition showed in general highest test-retest reliability as compared to all other measures.

### Results of voxel-wise reliability analyses

In the pooled analysis of both visits, robust task-evoked (de)activation was observed in tb-fMRI that is consistent with previous reports on these tasks for all paradigms except the Go/No-go (Fig 1). For the Go/No-go task, significant activation was only observed in the contrast Go>No-go in primary motor and insular regions but not in the opposite contrast. Activation patterns obtained for all tasks are shown separately for visits 1 and 2 in Figs S7–S13 at an uncorrected alpha level threshold of *p*<0.001 (except the Go/No-Go task, for which no significant activation was found when visits 1 and 2 were analyzed independently).

**Fig 1.**
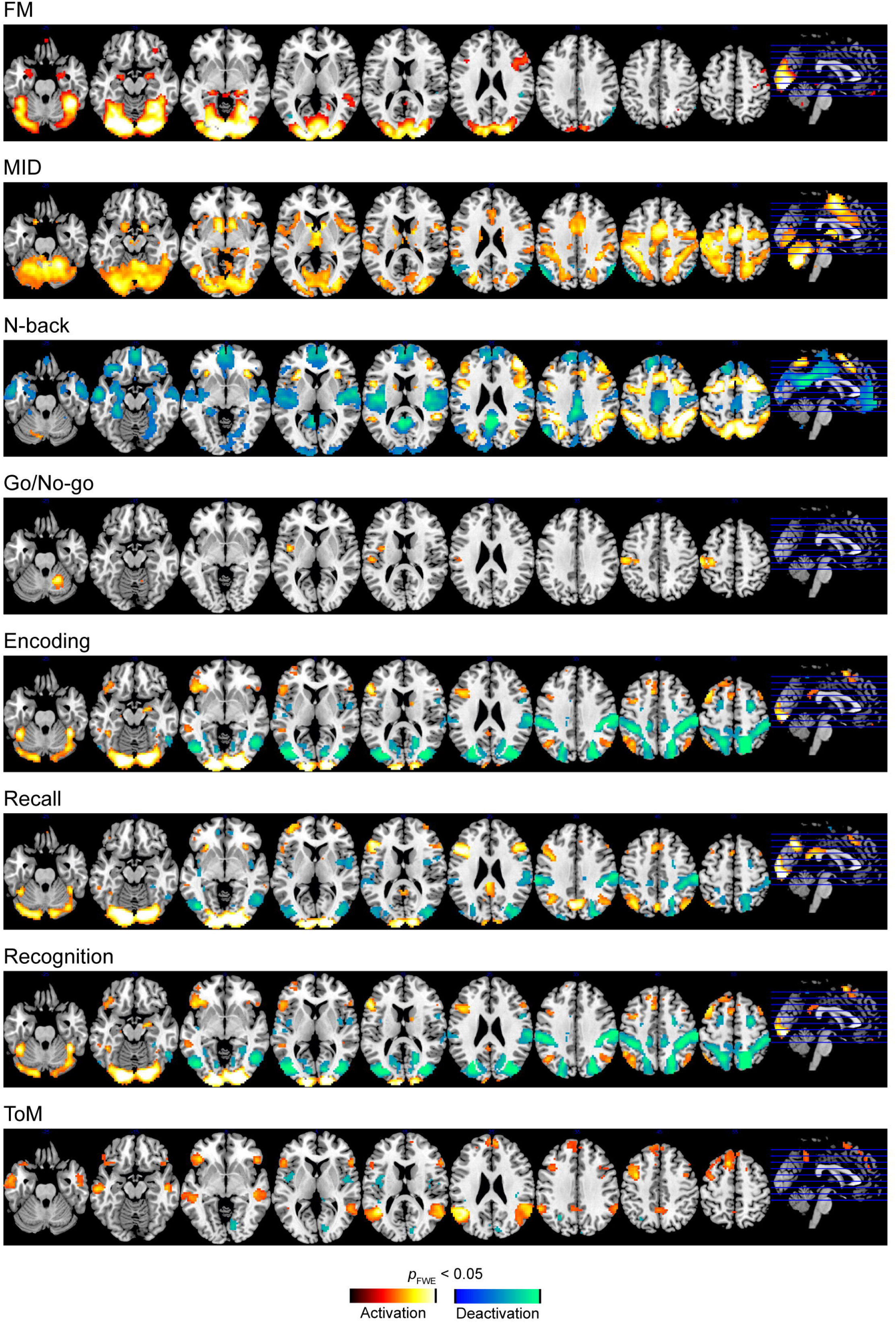
Group-level maps of main effects of all tb-fMRI responses. FWE: family-wise error corrected, MID: monetary incentive delay, tb-fMRI: task-based fMRI, ToM: theory of mind.

Jaccard indices of overlap between tb-fMRI activation patterns obtained at visit 1 and 2 revealed a generally high activation consistency for all tasks except Go/No-go (Table 1, Fig 2a). Highest consistency was achieved at a *t*-value threshold of 0 in all tasks except FM. Taken together, these results suggest overall consistency of activation vs. de-activation of fMRI patterns, even at very low significance thresholds. In the FM task, the highest consistency was achieved at a high positive *t*-value. Higher ICCs were observed in activated compared to deactivated regions, ranging between poor and good depending on the paradigm (Table 2, Fig 3). Evaluation of spatial reliability of average group activation maps obtained at visits 1 and 2 revealed a generally good (Go/No-go) to excellent (all other tasks) reliability of these tb-fMRI measures (Table 2). Finally, we found a significant positive correlation between the number of significantly activated or de-activated regions in the fMRI tasks and the observed test-retest reliability in the respective regions (*r* = 0.65; *p* = 0.008; Fig 2b).

**Table 1.** Consistency of tb-fMRI contrasts

| fMRI task | Number of de / activated voxels* at |  |  | Consistency |
| --- | --- | --- | --- | --- |
|  | Visit 1 | Visit 2 | Both | Jaccard index /<br>t-value threshold |
| **MID** | 15 / 893 | 222 / 1791 | 510 / 13551 | 0.900 / 0.01 |
| **N-back** | 4579 / 1839 | 2037 / 2306 | 10917 / 5573 | 0.871 / -0.10 |
| **ToM** | 61 / 919 | 24 / 248 | 475 / 3856 | 0.719 / 0.01 |
| **FM** | 20 / 3983 | 0 / 4427 | 122 / 6331 | 0.81 / 5.61 |
| **Encoding** | 3727 / 821 | 866 / 1697 | 5852 / 3848 | 0.695 / -0.09 |
| **Recall** | 1188 / 1519 | 335 / 1998 | 3364 / 4465 | 0.643 / -0.06 |
| **Recognition** | 157 / 1990 | 88 / 1962 | 1220 / 6229 | 0.769 / 0.01 |
| **Go/no-go** | 0 / 24 | 0 / 208 | 0 / 568 | 0.429 / 0.01 |

\* FWE-corrected significance at voxel-level ( $p < .05$ ), FM: face matching, MID: monetary incentive delay, ToM: theory of mind.

**Fig 2.**
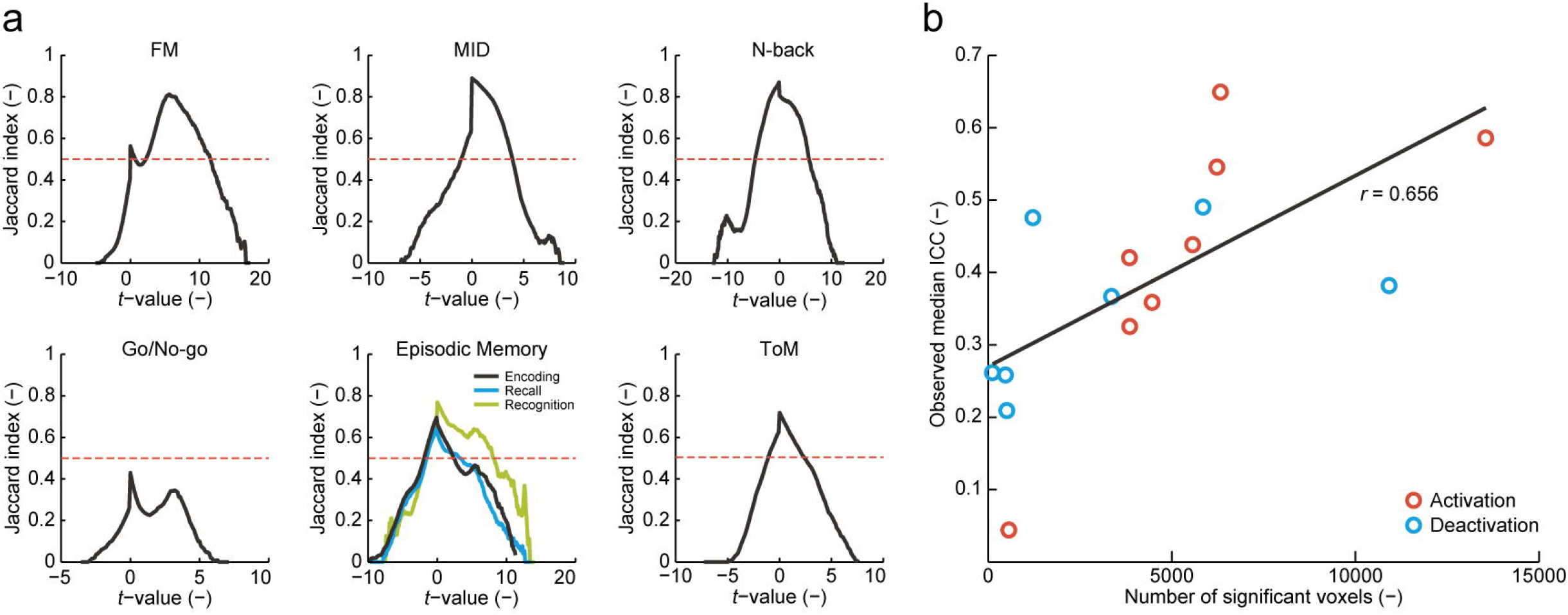
Consistency of (de)activation patterns (Jaccard index of overlap) (a) and association between number of significant voxels and observed reliability estimates observed for each fMRI task (b). FM: face matching, ICC: intraclass correlation coefficient, MID: monetary incentive delay, ToM: theory of mind.

**Fig 3.**
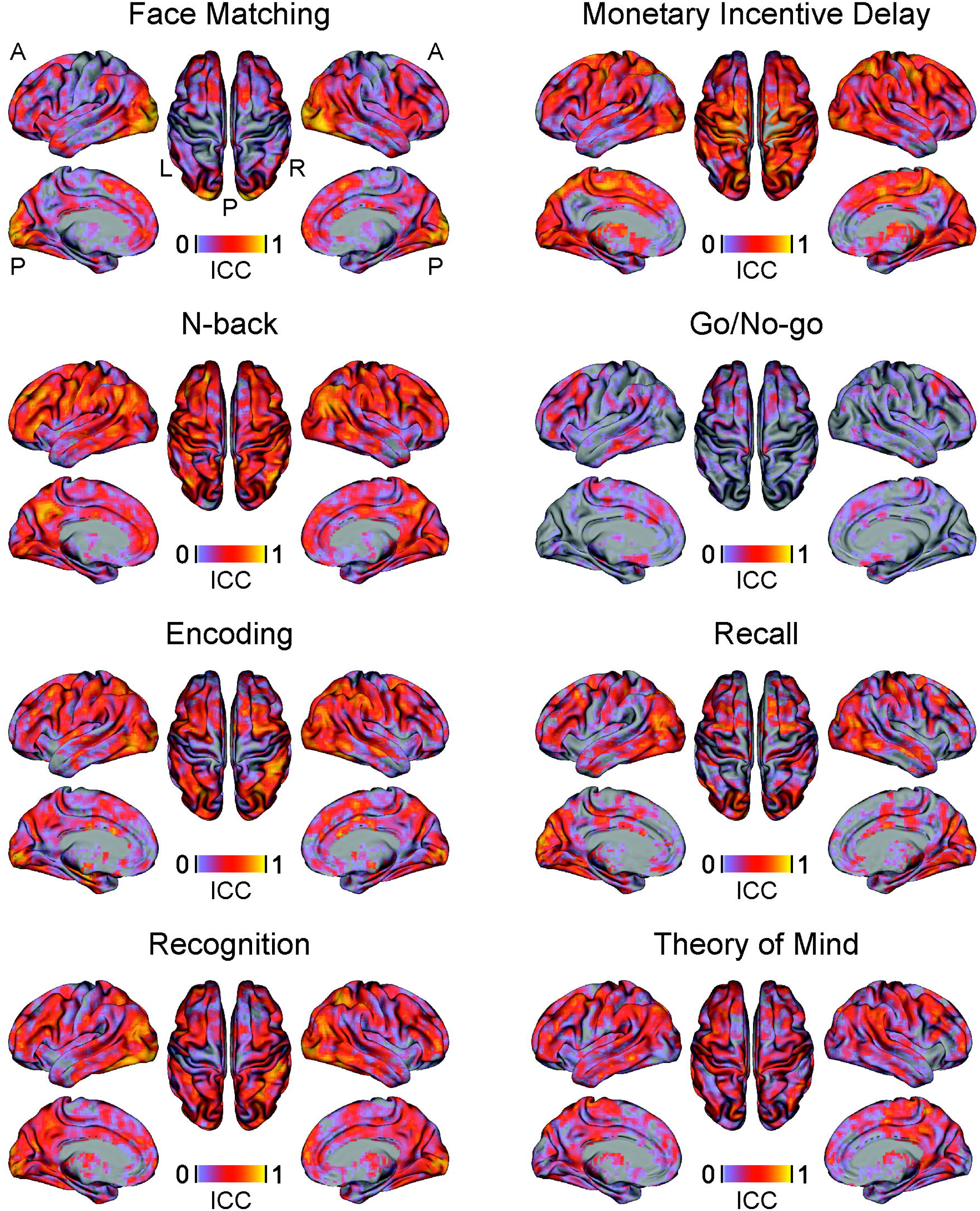
Voxel-wise reliability of tb-fMRI responses. ICC: intraclass correlation coefficient, tb-fMRI: task-based fMRI.

Voxel-wise and whole-brain ICC analyses of rs-fMRI data revealed a poor to excellent reliability for the different rs-fMRI measures depending on the pre-specified network (Table 3, Table S4, Fig 4). In general, lower ICCs and poor reliability estimates were obtained for whole-brain hubness or signal complexity measures (DC, EC and Hurst) compared to local activity and synchronization measures (ALFF, fALFF and ReHo). The reliabilities between rs-fMRI ICCs between screening and visit 1 and between visits 1 and 2 were comparable. ICCs for CBF ranged between fair and excellent with substantial ICC increases observed between visits 1 and 2 compared to between screening and visit 1 (Fig 4, Table 3, Table S4).

**Fig 4.**
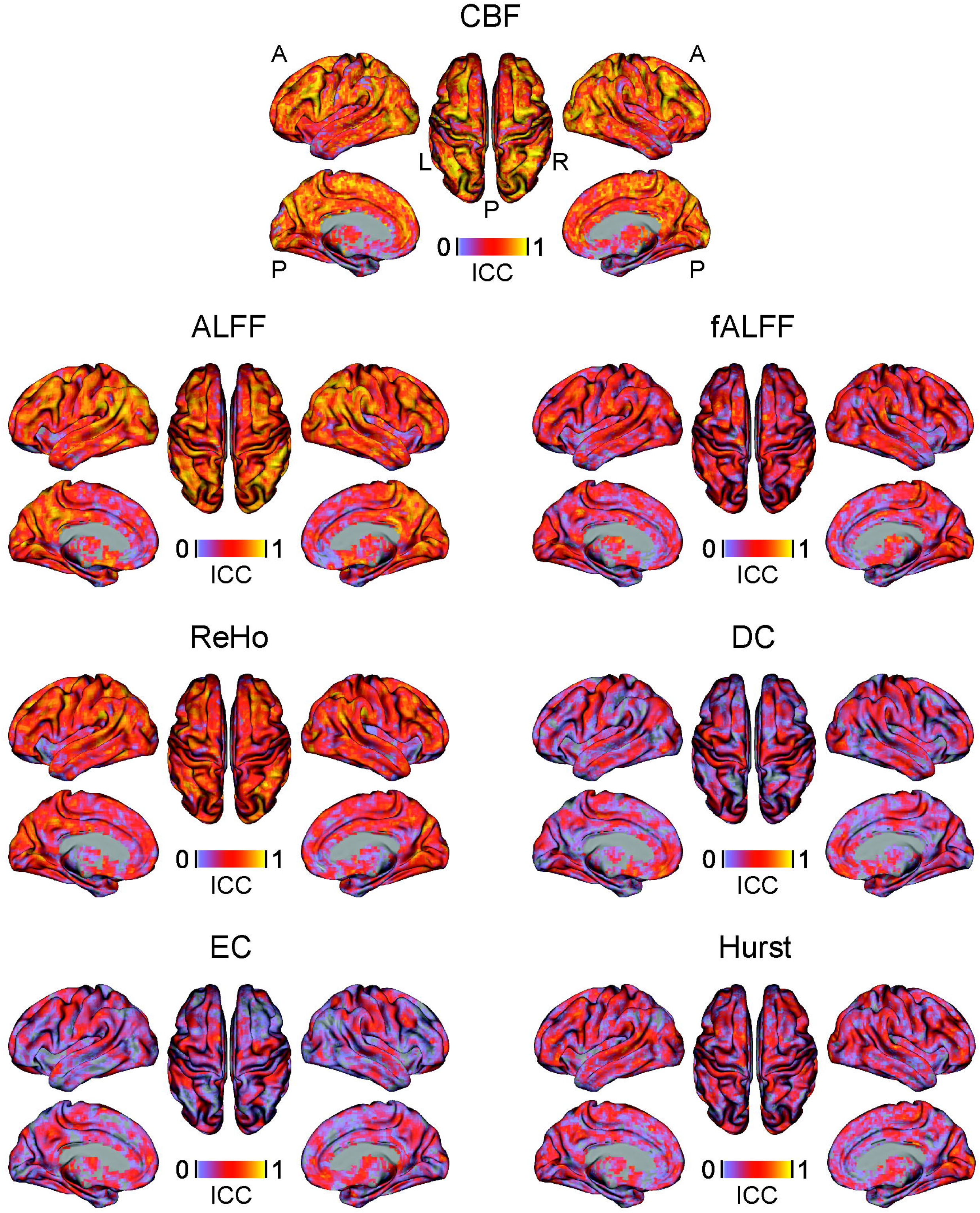
Voxel-wise reliability of rs-fMRI and CBF measures. ALFF: amplitude of low frequency fluctuations, fALFF: fractional ALFF, CBF: cerebral blood flow, DC: degree centrality, EC: eigenvector centrality, ICC: intraclass correlation coefficient, ReHo: regional homogeneity, rs-fMRI: resting-state fMRI.

### Results of region-wise reliability analyses

In tb-fMRI measures, the direction and magnitude of changes in ICCs from voxel-to region-wise analyses strongly depended on the specific paradigm and the regions chosen (Table 1, 2). Task specific ROI analyses revealed excellent test-retest reliability for MID and ToM and poor reliabilities for all other tasks (Table S3).

**Table 2.** Reliability of tb-fMRI contrasts

| fMRI task | Voxel-wise reliability<sup>2</sup> within |  | Reliability<sup>1</sup> of group t-maps | Reliability<sup>1</sup> of mean |  |
| --- | --- | --- | --- | --- | --- |
| | Deactivated regions**<br>median [$P_5$–$P_{95}$] | Activated regions**<br>median [$P_5$–$P_{95}$] | ICC [95% CI] | Deactivation**<br>ICC [95% CI] | Activation**<br>ICC [95% CI] |
| **MID** | 0.21 [-0.11–0.5] | 0.59 [0.25–0.78] | 0.93 [0.93–0.94] | -0.07 [-1.24–0.49] | 0.88 [0.75–0.94] |
| **N-back** | 0.38 [0.01–0.66] | 0.44 [0.13–0.68] | 0.97 [0.97–0.97] | 0.42 [-0.23–0.73] | -0.21 [-1.58–0.43] |
| **ToM** | 0.26 [-0.21–0.53] | 0.33 [-0.03–0.60] | 0.9 [0.9–0.9] | 0.51 [-0.02–0.77] | 0.52 [-0.01–0.77] |
| **FM** | 0.26 [-0.12–0.61] | 0.65 [0.13–0.87] | 0.95 [0.95–0.95] | 0.37 [-0.33–0.70] | 0.53 [0.01–0.78] |
| **Encoding** | 0.49 [0.05–0.76] | 0.42 [-0.01–0.77] | 0.94 [0.94–0.95] | 0.35 [-0.36–0.69] | 0.09 [-0.92–0.56] |
| **Recall** | 0.37 [-0.10–0.67] | 0.36 [-0.08–0.70] | 0.94 [0.93–0.94] | 0.42 [-0.22–0.72] | 0.22 [-0.63–0.63] |
| **Recognition** | 0.48 [0.07–0.75] | 0.55 [0.17–0.79] | 0.95 [0.95–0.95] | 0.66 [0.28–0.84] | 0.66 [0.29–0.84] |
| **Go/no-go** | – | 0.04 [-0.21–0.30] | 0.71 [0.70–0.71] | – | -0.71 [-2.6–0.18] |

\*\* FWE-corrected significance at voxel-level ( $p < .05$ ) in the pooled analyses of visit 1 and visit 2 data, <sup>1</sup> ICC(2,k), <sup>2</sup> ICC(2,1), CI: confidence interval, FM: face matching, ICC: intraclass correlation coefficient, MID: monetary incentive delay, $P$ : percentile, ToM: theory of mind.

In general, region-wise AAL-based analyses improved the reliability of rs-fMRI and CBF measures to a fair to excellent level (Table 3). Similar to voxel-wise analyses, a substantial increase in reliability from screening to visit 1 as compared to visit 1 to 2 was observed only for CBF..

**Table 3.**
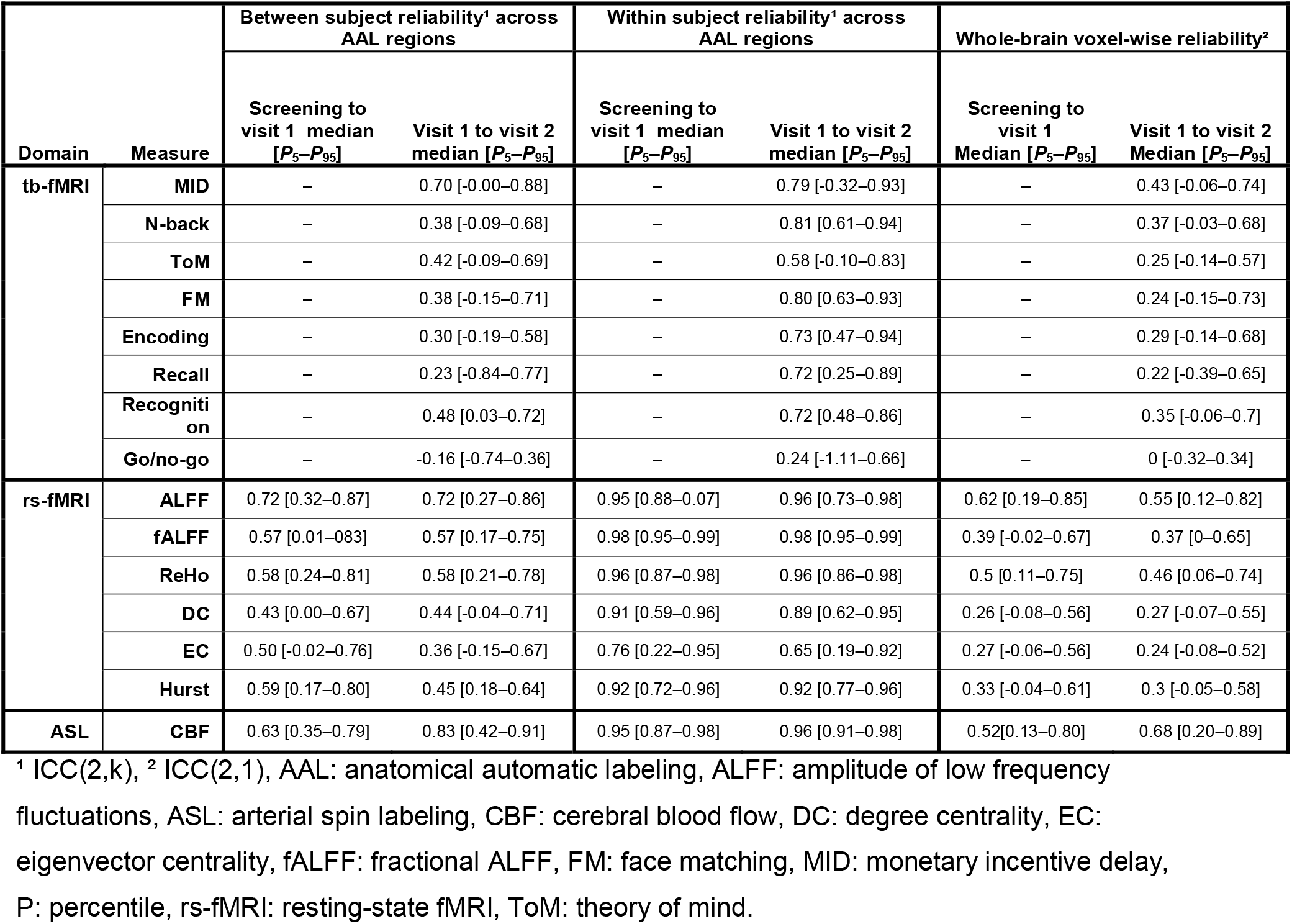
Comparative reliability of tb-fMRI, rs-fMRI and CBF measures

For all tb-fMRI, rs-fMRI and CBF measures, generally higher ICCs were observed for within-subject between-region compared to between-subject within-region ICCs, ranging from poor for Go/No-go, fair for ToM and good to excellent for all other measures (Table 3).

## Discussion

We evaluated different test-retest reliability characteristics for multiple MRI measures including tb-fMRI, rs-fMRI and CBF. We find a large heterogeneity of reliability estimates within and between the different domains, depending on the respective fMRI domain, analysis approach, reliability metric and study design.

Consistent with previous studies, voxel-wise test-retest reliability within (de)activated regions ranged between poor and good depending on the fMRI paradigm [20,23]. Also, similarly to previous studies, we obtained an excellent test-retest reliability of voxel-wise group activation maps for all fMRI tasks [23]. No consistent improvement in test-retest reliability was observed after averaging signals from the (de)activated regions: some tasks showed substantial improvements but also considerable worsening in respective reliability metrics. Interestingly, we found that the amount of significant activation (i.e., number of activated voxels) was positively related to the task and region-specific reliability estimates. This observation suggests that differences in amount of activation may be used as a first reliability estimate of novel tb-fMRI paradigms. Furthermore, we found strong differences between fMRI tasks with respect to consistency of between-session activation patterns and the respective dependence on the applied statistical thresholds. Interestingly, for most tasks, highest consistency of between-session activation maps was achieved at a zero threshold, suggesting that whole brain (de)activation patterns are reliable despite most regions not reaching the significance threshold. In contrast, for the face matching task, highest consistency was observed at a relatively large *t*-value, indicating a highly reliable activation pattern for this task. For most other fMRI tasks, the consistency of activation patterns significantly dropped at such high thresholds, indicating a rather low regional overlap of peak activations across different visits.

Relative to tb-fMRI, higher between-subject test-retest reliability was observed in rs-fMRI local activity measures (ALFF, fALFF and ReHo), which ranged from good to excellent. In contrast, global connectivity (DC and EC) and signal complexity measures (Hurst) showed poor to fair test-retest reliabilities. The large heterogeneity of reliability estimates across different rs-fMRI metrics is consistent with previous reports [17,36]. However, the present findings suggest that especially voxel-wise connectivity metrics provide poor between-subject reliability in a healthy volunteer setting. In contrast, the within-subject between-region reliability of all rs-fMRI measures was in the excellent range and consistently higher than the between-subject reliability, indicating a high topographic stability of all rs-fMRI measures. The within-subject ICC may be considered an indicator of the amount of information carried by the respective measure. In contrast, as the between-subject ICC reflects regional signal-to-noise levels in the respective measure, lower values observed here suggest that these measures may be insufficient for correlational analyses (e.g. with behavioral scales). Overall poor performance of the evaluated connectivity-based rs-fMRI metrics questions their usability for cross-sectional correlation analyses in healthy volunteers.

Several recent studies established the test-retest reliability of CBF and connectivity measures and the potential influence of acquisition and preprocessing parameters, but also effects of different approaches for calculating a particular outcome measure [16,18,24,25,37–40]. Consistent with these reports, we observed excellent test-retest reliability for quantitative CBF, which outperformed most rs-fMRI and all tb-fMRI metrics [18,24]. As for rs-fMRI, ROI test-retest estimates were superior to voxel-based estimates. Additionally, we found a substantial improvement of test-retest characteristics between data from visits 1 and 2 as compared to screening and visit 1, but only for CBF and none of the rs-fMRI measures. A potential explanation for this pattern might be the stronger susceptibility of CBF to peripheral or central arousal effects [41,42]. Emotional arousal may be heightened in a first compared to ensuing MRI sessions, potentially introducing additional noise into the baseline CBF measures and thereby lowering its test-retest reliability with subsequent sessions. In contrast, the low frequency band-pass filtering and other de-noising techniques applied to rs-fMRI measures should reduce the contribution of such physiological confounds and might therefore explain the lack of changes in rs-fMRI reliability.

Several limitations apply to the interpretation of specific outcomes. As the major purpose of the present study was to compare the relative test-retest reliabilities of specific tb-fMRI, rs-fMRI and CBF, we used the recommended and most comparable pre-processing pipelines for these data. Additional pre-processing steps may therefore have further improved the reliability of some of the evaluated metrics (i.e. slice timing or different motion correction for some of the fMRI tasks). Same issue also applies to the choice of parameter settings such as the correlational threshold applied for DC in our study. The choice of different thresholds has been shown to affect functional connectivity results and may therefore also result in different test-reliability estimates [43]. Similarly, we applied a conservative approach to correct for potential motion effects and white matter and cerebrospinal signals as recommended for rs-fMRI data [29], compared to the standard fMRI motion correction using 6 parameters. Such differences may have introduced further biases between the respective fMRI domains. However, considering that the observed ICCs for both fMRI and rsMRI ranged from poor to excellent, the effect of these differential processing steps may be negligible. Lastly, further differences between tb-fMRI, rs-fMRI and CBF measures may have been introduced through different acquisition parameters (i.e. lower resolution for CBF which have lowered its test-retest reliability estimates) and the study design with only rs-fMRI and CBF measures acquired at the screening visit.

To our knowledge, this study provides the most comprehensive evaluation of test-retest metrics for commonly used tb-fMRI and rs-fMRI measures. We find most of the rs-fMRI measures to have superior reliability compared to tb-fMRI. The relative reliabilities of fMRI measures strongly depended on the task, with more widespread activation associated with higher test-retest reliability. Lastly, we find the reliability of CBF to substantially benefit from an additional screening MRI evaluation, which may reduce potential emotional arousal effects and respective cardiovascular changes confounding the baseline CBF scan. Importantly, previous studies have demonstrated the dependency of achieved power to detect specific effects in both within- and between-subject designs on the test-retest reliability of respective metrics [44]. Our study provides an overview of different test-retest reliability metrics for the most commonly applied functional task-based and resting-state MRI domains and specific outcome measures. It therefore enables a more informed decision on end-point selection, study design, and sample size required to detect specific effect sizes with the respective technologies.

## Acknowledgements

We thank Anita Sibeijn-Kuiper and Judith Streurman from the Neuroimaging Center at University Medical Center Groningen for performing the MRI examinations and for their technical support and Kirsten Taylor for her copy-editing support.

## Declaration of interests

SH, FS, CL, GG, NS SAS, AB and JD are current or former full-time employees of F.Hoffmann-La Roche, Basel Switzerland. The authors received no specific funding for this work. F.Hoffmann-La Roche provided financial contribution in the form of salary for all authors but did not have any additional role in the study design, data collection and analysis, decision to publish, or preparation of the manuscript.

## Authors contributions

SH, FS, GG, NS, SAS, AB and JD designed the study, RJR and JCM conducted the study, SH, NS and JD performed the analyses. SH and JD wrote the manuscripts. All authors reviewed the manuscript and provided feedback.

## Supporting information

2017_Test-Retest_Supplement.docx

Data1.zip

Data2.zip

Data3.zip

## References

1. Fransson P. Spontaneous low-frequency BOLD signal fluctuations: An fMRI investigation of the resting-state default mode of brain function hypothesis. Hum Brain Mapp. 2005;26: 15–29.

2. Friston KJ, Fletcher P, Josephs O, Holmes A, Rugg MD, Turner R. Event-related fMRI: characterizing differential responses. Neuroimage. 1998;7: 30–40.

3. Kim J, Whyte J, Wang J, Rao H, Tang KZ, Detre JA. Continuous ASL perfusion fMRI investigation of higher cognition: quantification of tonic CBF changes during sustained attention and working memory tasks. Neuroimage. 2006;31: 376–385.

4. Lee MH, Smyser CD, Shimony JS. Resting-state fMRI: a review of methods and clinical applications. Am J Neuroradiol. 2013;34: 1866–1872.

5. Logothetis NK. What we can do and what we cannot do with fMRI. Nature. 2008;453: 869–878.

6. Newberg AB, Wang J, Rao H, Swanson RL, Wintering N, Karp JS, et al. Concurrent CBF and CMRGlc changes during human brain activation by combined fMRI-PET scanning. Neuroimage. 2005;28: 500–6.

7. Van Den Heuvel MP, Hulshoff Pol HE. Exploring the brain network: a review on resting-state fMRI functional connectivity. Eur Neuropsychopharmacol. 2010;20: 519–534.

8. Borsook D, Becerra L, Hargreaves R. A role for fMRI in optimizing CNS drug development. Nat Rev Drug Discov. 2006;5: 411–425. doi:10.1038/nrd2027

9. Chen Y, Parrish TB. Caffeine dose effect on activation-induced BOLD and CBF responses. NeuroImage. 2009;46: 577–583. doi:10.1016/j.neuroimage.2009.03.012

10. Dennis NA, Kim H, Cabeza R. Effects of aging on true and false memory formation: An fMRI study. Neuropsychologia. 2007;45: 3157–3166.

11. Handley R, Zelaya FO, Reinders AATS, Marques TR, Mehta MA, O’Gorman R, et al. Acute effects of single-dose aripiprazole and haloperidol on resting cerebral blood flow (rCBF) in the human brain. Hum Brain Mapp. 2013;34: 272–282. doi:10.1002/hbm.21436

12. Bruns A, Mueggler T, Künnecke B, Risterucci C, Prinssen EP, Wettstein JG, et al. “Domain gauges”: A reference system for multivariate profiling of brain fMRI activation patterns induced by psychoactive drugs in rats. NeuroImage. 2015;112: 70–85. doi:10.1016/j.neuroimage.2015.02.032

13. Koch W, Teipel S, Mueller S, Benninghoff J, Wagner M, Bokde AL, et al. Diagnostic power of default mode network resting state fMRI in the detection of Alzheimer’s disease. Neurobiol Aging. 2012;33: 466–478.

14. Matthews PM, Honey GD, Bullmore ET. Applications of fMRI in translational medicine and clinical practice. Nat Rev Neurosci. 2006;7: 732–744. doi:10.1038/nrn1929

15. Song X-W, Dong Z-Y, Long X-Y, Li S-F, Zuo X-N, Zhu C-Z, et al. REST: a toolkit for resting-state functional magnetic resonance imaging data processing. PloS One. 2011;6: e25031.

16. Jann K, Gee DG, Kilroy E, Schwab S, Smith RX, Cannon TD, et al. Functional connectivity in BOLD and CBF data: similarity and reliability of resting brain networks. NeuroImage. 2015;106: 111–122. doi:10.1016/j.neuroimage.2014.11.028

17. Braun U, Plichta MM, Esslinger C, Sauer C, Haddad L, Grimm O, et al. Test–retest reliability of resting-state connectivity network characteristics using fMRI and graph theoretical measures. Neuroimage. 2012;59: 1404–1412.

18. Chen Y, Wang DJ, Detre JA. Test–retest reliability of arterial spin labeling with common labeling strategies. J Magn Reson Imaging. 2011;33: 940–949.

19. De Simoni S, Schwarz AJ, O’Daly OG, Marquand AF, Brittain C, Gonzales C, et al. Test–retest reliability of the BOLD pharmacological MRI response to ketamine in healthy volunteers. NeuroImage. 2013;64: 75–90. doi:10.1016/j.neuroimage.2012.09.037

20. Friedman L, Stern H, Brown GG, Mathalon DH, Turner J, Glover GH, et al. Test–retest and between-site reliability in a multicenter fMRI study. Hum Brain Mapp. 2008;29: 958–972.

21. Klomp A, van Wingen GA, de Ruiter MB, Caan MWA, Denys D, Reneman L. Test–retest reliability of task-related pharmacological MRI with a single-dose oral citalopram challenge. NeuroImage. 2013;75: 108–116. doi:10.1016/j.neuroimage.2013.03.002

22. Manoach DS, Halpern EF, Kramer TS, Chang Y, Goff DC, Rauch SL, et al. Test-retest reliability of a functional MRI working memory paradigm in normal and schizophrenic subjects. Am J Psychiatry. 2001;158: 955–958.

23. Plichta MM, Schwarz AJ, Grimm O, Morgen K, Mier D, Haddad L, et al. Test–retest reliability of evoked BOLD signals from a cognitive–emotive fMRI test battery. Neuroimage. 2012;60: 1746–1758.

24. Fazlollahi A, Bourgeat P, Liang X, Meriaudeau F, Connelly A, Salvado O, et al. Reproducibility of multiphase pseudo-continuous arterial spin labeling and the effect of post-processing analysis methods. NeuroImage. 2015;117: 191–201. doi:10.1016/j.neuroimage.2015.05.048

25. Zou Q, Miao X, Liu D, Wang DJJ, Zhuo Y, Gao J-H. Reliability comparison of spontaneous brain activities between BOLD and CBF contrasts in eyes-open and eyes-closed resting states. NeuroImage. 2015;121: 91–105. doi:10.1016/j.neuroimage.2015.07.044

26. Williams EJ. Experimental designs balanced for the estimation of residual effects of treatments. Aust J Chem. 1949;2: 149–168.

27. Alsop DC, Detre JA, Golay X, Günther M, Hendrikse J, Hernandez-Garcia L, et al. Recommended implementation of arterial spin-labeled perfusion MRI for clinical applications: A consensus of the ISMRM perfusion study group and the European consortium for ASL in dementia. Magn Reson Med. 2015;73: 102–116.

28. Friston KJ, Williams S, Howard R, Frackowiak RS, Turner R. Movement-related effects in fMRI time-series. Magn Reson Med. 1996;35: 346–355.

29. Yan C-G, Craddock RC, He Y, Milham MP. Addressing head motion dependencies for small-world topologies in functional connectomics. Front Hum Neurosci. 2013;7.

30. Wink AM, de Munck JC, van der Werf YD, van den Heuvel OA, Barkhof F. Fast eigenvector centrality mapping of voxel-wise connectivity in functional magnetic resonance imaging: implementation, validation, and interpretation. Brain Connect. 2012;2: 265–274.

31. Maxim V, Şendur L, Fadili J, Suckling J, Gould R, Howard R, et al. Fractional Gaussian noise, functional MRI and Alzheimer’s disease. Neuroimage. 2005;25: 141–158.

32. Buckner RL, Sepulcre J, Talukdar T, Krienen FM, Liu H, Hedden T, et al. Cortical hubs revealed by intrinsic functional connectivity: mapping, assessment of stability, and relation to Alzheimer’s disease. J Neurosci. 2009;29: 1860–1873.

33. Weir JP. Quantifying test-retest reliability using the intraclass correlation coefficient and the SEM. J Strength Cond Res. 2005;19: 231–240.

34. Cicchetti DV, Sparrow SA. Developing criteria for establishing interrater reliability of specific items: applications to assessment of adaptive behavior. Am J Ment Defic. 1981; Available: http://psycnet.apa.org/psycinfo/1982-00095-001

35. McGraw KO, Wong SP. Forming inferences about some intraclass correlation coefficients. Psychol Methods. 1996;1: 30.

36. Telesford QK, Morgan AR, Hayasaka S, Simpson SL, Barret W, Kraft RA, et al. Reproducibility of graph metrics in fMRI networks. Front Neuroinformatics. 2010;4: 117.

37. Liang X, Wang J, Yan C, Shu N, Xu K, Gong G, et al. Effects of different correlation metrics and preprocessing factors on small-world brain functional networks: a resting-state functional MRI study. PloS One. 2012;7: e32766.

38. Schwarz AJ, McGonigle J. Negative edges and soft thresholding in complex network analysis of resting state functional connectivity data. Neuroimage. 2011;55: 1132–1146.

39. Tancredi FB, Lajoie I, Hoge RD. Test-retest reliability of cerebral blood flow and blood oxygenation level-dependent responses to hypercapnia and hyperoxia using dual-echo pseudo-continuous arterial spin labeling and step changes in the fractional composition of inspired gases. J Magn Reson Imaging JMRI. 2015;42: 1144–1157. doi:10.1002/jmri.24878

40. Wang J-H, Zuo X-N, Gohel S, Milham MP, Biswal BB, He Y. Graph theoretical analysis of functional brain networks: test-retest evaluation on short-and long-term resting-state functional MRI data. PloS One. 2011;6: e21976.

41. Gianaros PJ, Van der Veen FM, Jennings JR. Regional cerebral blood flow correlates with heart period and high-frequency heart period variability during working-memory tasks: Implications for the cortical and subcortical regulation of cardiac autonomic activity. Psychophysiology. 2004;41: 521–530.

42. Wang J, Rao H, Wetmore GS, Furlan PM, Korczykowski M, Dinges DF, et al. Perfusion functional MRI reveals cerebral blood flow pattern under psychological stress. Proc Natl Acad Sci U S A. 2005;102: 17804–17809.

43. Wijk BCM van, Stam CJ, Daffertshofer A. Comparing Brain Networks of Different Size and Connectivity Density Using Graph Theory. PLOS ONE. 2010;5: e13701. doi:10.1371/journal.pone.0013701

44. Kanyongo GY, Brook GP, Kyei-Blankson L, Gocmen G. Reliability and statistical power: How measurement fallibility affects power and required sample sizes for several parametric and nonparametric statistics. J Mod Appl Stat Methods. 2007;6: 9.

